# Context-dependent effects of microglial MyD88 removal on voluntary ethanol consumption in mice

**DOI:** 10.64898/2026.06.12.731650

**Authors:** Julia E. Dziabis, Neil Rogers, Benjamin L. Horvath, Michael S. Patton, Irene O. Jonathan, Erica J. Freeman, Will Sun, Jerome Moulden, Grace Zhang, Staci D. Bilbo

## Abstract

Neuroimmune signaling is increasingly implicated in alcohol use disorder (AUD). Microglia, the brain’s resident immune cells, signal in part through the adaptor protein myeloid differentiation primary response 88 (MyD88), a key mediator of innate immune responses. Here, we investigated whether microglial-specific MyD88 signaling regulates voluntary alcohol consumption in adulthood, as whole-body loss of MyD88 was previously shown to increase drinking. We further determined if alcohol altered parvalbumin-expressing interneurons (PVIs) and microglia within the pre-frontal cortex, based on our previously described role for MyD88 signaling on perineuronal net (PNN) deposition on PVIs in several brain regions, and the well characterized role of inhibitory signaling in alcohol use disorders.

Loss of microglial-MyD88 had minimal effects on voluntary alcohol intake and anxiety-like behaviors. Alcohol exposure did not modify observed MyD88-dependent changes in PVIs/PNNs, despite altering microglial morphology in the male prefrontal cortex independent of genotype. The addition of an early life endotoxin challenge was sufficient to induce an increase in adult alcohol consumption in both MyD88-deficient and control males. However, injection of saline alone also induced an increase in adult drinking in MyD88-deficient males.

These findings suggest that microglial-MyD88 signaling does not strongly regulate alcohol intake under baseline conditions in a one-bottle, voluntary binge-drinking paradigm, however there may be a role for microglial-MyD88 signaling in modulating the impact of developmental environmental contexts, such as stress, in later-life male drinking behavior. This work highlights the importance of developmental context, such as stress or inflammatory history, in understanding underlying microglia signaling mechanisms in conferring AUD risk.

## 1. Introduction

Heavy alcohol consumption is the main risk factor for alcohol use disorder (AUD), a devastating disease characterized by an inability to stop drinking despite causing harm to oneself and others (Kranzler and Soyka, 2018). Binge drinking and heavy alcohol use are a massive public health burden and have increased in the United States, particularly in the last few years, during and following the COVID-19 pandemic (Grossman et al., 2020). Despite advances in our understanding of brain changes in AUD, existing pharmacological treatments are limited in their effectiveness. Understanding the neurobiological factors important for initial excessive drinking behaviors, and the progression to AUD, could enhance both prevention and treatment of AUD.

Recent attention has shifted to the neuroimmune system as a potential therapeutic target for AUD. This interest is supported by studies in humans with AUD identifying gene expression adaptations to chronic drinking; unbiased gene expression analyses consistently revealed robust changes in immune function in the human frontal cortex (Lewohl et al., 2000; Liu et al., 2006; Mayfield et al., 2002). Work in rodents confirmed these changes and have attempted to identify the signaling pathways that are most critical for behavioral responses to alcohol, such as craving, consumption, tolerance, and dependence (Robinson et al., 2014). Alcohol likely acts through multiple innate immune toll-like receptors (TLRs) on microglia, both directly and indirectly, through increased release of reactive oxygen species and cytokines, to stimulate the release of immune signaling molecules from microglia and other immune cells within the brain. Microglia, the brain-resident macrophages, use these same neuroimmune molecules to interact closely with neurons and their synapses, providing a mechanism through which alcohol impacts neuroplasticity and behavior (Williamson and Bilbo, 2013; Wu et al., 2015).

Indeed, both in human post-mortem tissue and in rodent models of AUD, microglia are continually identified as a cell type of interest in trying to understand AUD pathology. While microglia are unsurprisingly acutely responsive to alcohol (Crews et al., 2017), the impact of alcohol on microglia and their long-term function is variable depending on the timing of drinking, the dose or amount consumed, and even the brain region observed (Sanchez-Alavez et al., 2019). In general, in rodents models of forced, non-acute alcohol exposure, such as through alcohol feeding, alcohol gavage or injection, or vaporized inhalation of alcohol, microglia exhibit dramatic changes in morphology suggestive of a pro-inflammatory state, often accompanied by increases in pro-inflammatory signaling when also quantified (Avila et al., 2017; Paouri et al., 2025; Qin and Crews, 2012a; Walter et al., 2017; Warden et al., 2020a; Zhao et al., 2013). These findings are in alignment with observations made in post-mortem brains of individuals that suffered from AUD, where microglia also display altered morphology, depending on the brain region sampled (Dennis et al., 2014; He and Crews, 2008; Rasool et al., 2024; Rubio-Araiz et al., 2017). While the functional consequences of this microglial response to alcohol exposure are still being investigated, recent evidence has supported the idea that microglia may be critical specifically for the transition between non-dependent (more mild) alcohol consumption and alcohol dependence in male mice (Warden et al., 2021, 2020b). Because these studies relied on total broad pharmacological depletion of microglia, elucidating the specific signaling pathways in microglia that are important in excessive binge drinking behaviors remains an interesting avenue of research to pursue.

Previous studies have sought to identify which aspects of TLR signaling pathways may be involved in excessive alcohol consumption. However, of the many studies in rodents removing different aspects of TLR signaling (e.g. TLR2-KO, TLR4-KO, etc.), only the global loss of MyD88 led to increased voluntary drinking behavior (Blednov et al., 2017). MyD88, or myeloid differentiation primary response 88, is an essential co-adaptor protein for nearly every TLR, meaning it has broad importance in a wide range of inflammatory responses. Interestingly, some TLRs have a MyD88-independent signaling pathway that leads to the upregulation of type I interferon genes, which are increased in the brains of human alcoholics and following excessive alcohol drinking in mice (Erickson et al., 2019; Kapoor et al., 2019). While primarily microglia express MyD88 in the brain, other cell types such as astrocytes and neurons also express MyD88. Therefore, definitively implicating microglial-MyD88 signaling in increased drinking behavior is of great relevance to the field in terms of understanding the brain-specific mechanisms of excessive consumption.

## 2. Methods

### 2.1 Animals

All experiments were conducted in accordance with the NIH *Guide to the Care and Use of Laboratory Animals* and approved by the Duke University Institutional Animal Care and Use Committee (IACUC). Animals were group housed in a standard 12-12h light/dark cycle. MyD88-floxed mice were obtained from Jackson Laboratory (Stock #008888) and crossed to Cx3cr1-BT-Cre (MW126GSat), which were provided by L. Kus (GENSAT BAC Transgenic Project) until all offspring were fully MyD88 floxed (F/F) and either Cre-negative (Cre^0/0^ or CON) or Cre-positive (Cre^tg/0^ or cKO). Mice were then backcrossed onto a fully Jackson background. The Cre transgene was maintained in males for all experimental studies whenever possible, and has minimal off-target effects in microglia (Mroue-Ruiz et al., 2026). Recombination efficiency in this line has been previously validated in brain tissue (Rawls et al., 2025), however, Cx3cr1-driven recombination is not restricted to brain microglia and may also affect the expression of MyD88 in Cx3cr1-expressing tissue-resident macrophages outside the brain. Each litter was bred to contain a mixture cKO offspring and CON offspring to be used as littermate controls. CON and cKO experimental mice were co-housed for all experiments. For all experimental endpoints, mice from at least 3 different litters were included in each sex/genotype/treatment group, unless otherwise noted.

For all drinking experiments in adults, mice were transferred to a room with a new light cycle schedule and allowed to habituate to the new room with water bottles instead of lixits for 2 weeks. All mice were single housed 1 week prior to the start of drinking paradigms. All water was provided by bottles rather than lixits to allow for exclusive access to only alcohol for the short drinking periods (2-4h), then water bottles were returned for *ad libitum* access. Mice always had access to food before, during, and after alcohol access.

### 2.2 Genotyping

Genotyping of transgenic animals was conducted from tail (at postnatal day (P)14) or toe (at P4) DNA using polymerase chain reaction (PCR). For all experimental animals, genotyping confirmed fully floxed MyD88 alleles as well as the absence or presence of Cre recombinase (Table 1). High recombination efficiency has been previously validated in this mouse line by PCR followed by gel electrophoresis (Rawls et al., 2025).

**Table 1:**
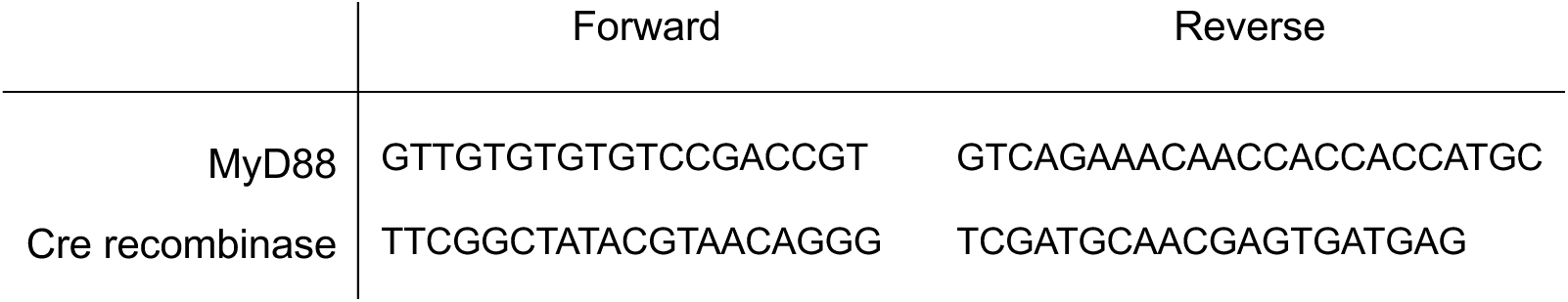
Genotyping Primer Sequences.

### 2.3 Adult Drinking in the Dark (DID) alcohol procedure

To quantify alcohol consumption, mice were single housed for 1 week prior to experiment start, and water access was provided through bottles with metal sippers to reduce novelty aversion to alcohol sippers. Mice were weighed at the start of each week, and then 3 hours into the “dark” cycle (lights off), the water bottle was switched with a modified syringe sipper containing 20% w/v ethanol in water for 2 hours. Ethanol was made fresh daily. Volume of alcohol consumed was recorded at the conclusion of the period and then normalized to each mouse’s body weight for that week (g/kg consumed). Days 1, 2, and 3 of each week were as above, and Day 4 was the same except with 4 hours of access (“binge day”). Water controls underwent the same exact procedures, except with water in modified sippers instead of alcohol. The bitterant quinine (0.3mM) was added to the same ethanol solution for quinine aversion-resistant drinking procedure. Ethanol + quinine was provided for 4 hours following the same procedure as the “binge day.”

### 2.4 Sucrose preference procedure

Sucrose preference was performed as previously described (Ceasrine et al., 2022). In brief, singly housed mice were provided with 2 bottles to drink from for 72 hours. One bottle contained normal drinking water, and the other contained 1% sucrose in drinking water. Intake from each bottle was measured by weight every 24 hours, and the position of bottles (left vs. right side of the cage) were rotated daily to reduce potential side preference bias. Sucrose preference was calculated as follows: ((sucrose consumption)/(sucrose + water consumption)) × 100, and then the 3 measurements were averaged for each animal.

### 2.5 Intermittent access 2-bottle choice (2-BC) procedure

Following 6 weeks of DID, mice underwent an intermittent ethanol exposure paradigm, a regimen consisting of access to 2 bottles in home cages, one filled with water and the other filled with 20% w/v ethanol. Access to ethanol followed a weekly repeating schedule of 1 day off, 1 day on, 1 day off, 1 day on, 2 day off, 1 day on in order to create intermittent access. During “on” access days, both ethanol and water bottles remained in the cage for 24 hours. After 24 hours, both bottles were removed from the cage, weighed, washed, and both refilled with water for the “off” access day. The location of the alcohol bottle versus the water bottle was alternated between the left and right side of the cage each “on” day to control for any side preference. Water access only controls were also assessed, with access to 2 water bottles each day.

### 2.6 Anxiety-like behavior assays

All behavior was run during the dark phase, around the same time that mice would normally receive alcohol (∼3h into dark cycle). Mice were kept away from light until the time of testing, where they experienced anxiety-like procedures in bright overhead light just for the duration of testing.

For elevated plus maze (EPM), mice were placed on a X-shaped apparatus, where one pair of arms are open, without walls, and the other two arms are “closed,” with high walls providing shelter. Rodents are prey animals and therefore have an evolutionary drive to avoid open spaces, such as the open arms of the EPM. Mice were placed at one end of one the closed arms, and behavior recorded for a 5-minute period from an overhead camera. Ethovision was used to quantify the time spent in open vs. closed arms, with an open arm entry counted when all four limbs were outside of the center of the maze and on the open arm.

Light-dark (L-D) box was performed with a square shaped apparatus, where half of the square is enclosed with black opaque walls and a lid, and a small opening that connects to the other, open half of the square. Mice were placed in the dark side of the apparatus, the lid was placed, and the opening connecting the two halves was covered. Trial began when the doorway connecting the two halves was uncovered. Mice were recorded freely exploring for 10 minutes using an overhead camera, and time spent in on either side of the testing apparatus was quantified using Ethovision. Entry into the open side of the chamber was quantified as all four limbs exiting the closed chamber.

### 2.7 Immunohistochemistry

Mice were sacrificed by CO_2_ inhalation. Mice were then transcardially perfused with ice-cold saline followed by 4% paraformaldehyde (PFA) and brains post-fixed in 4% PFA for 24 hours at 4°C. Brains were then dehydrated in a 30% sucrose in 1X PBS solution with 0.1% sodium azide for at least 72 hours at 4°C. Brains were flash-frozen in 2-methylbutane pre-chilled on dry-ice and stored at −80°C until cryosectioning at 40μm into tubes of cryoprotectant. Tissue was stored at −20°C until use.

For all endpoints, free floating tissue sections were washed 3 times in 1X PBS at room temperature (RT) on a shaker for 10 minutes, then blocked in 5% or 10% normal goat serum (NGS) and 0.1 or 0.3% Triton-X in 1X PBS for 1-2 hours at RT on a shaker. Tissue was then moved into primary solution, which matched blocking solution with the addition of primary antibodies (Rabbit α Parvalbumin, Swant #PV27a, 1:1000; Biotin WFA, Vector Laboratories #B-1355, 1:1500; Chicken α Iba1, Synaptic Systems #234009, 1:1000; 1° Guinea Pig α Tmem119; Synaptic Systems #400004, 1:500; Rat α CD68, Biolegend #137002, 1:500), for an overnight incubation at 4°C. Primary solution was then washed off by washing 3 times in 1X PBS at RT on the shaker for 10 minutes. Tissue was incubated, covered, on a shaker at RT for 2-4 hours with corresponding secondary antibodies in 1X PBS (ThermoFisher, 1:500), then washed 3 times again before mounting on subbed slides and coverslipping with Vectashield Plus with DAPI.

### 2.8 Adult interneuron and extracellular matrix quantification

Immunohistochemistry was performed as described above with appropriate antibodies. Tilescans of the prefrontal cortex, striatum, and dorsal hippocampus, were captured on a Zeiss Axio Imager Z1 epifluorescent microscope with apotome attachment using a 10X objective. Maximum intensity projections were produced from 19-step Z stacks captured with a 1.67μm step size. Projected images were imported into QuPath for manual blinded counting of PV cells and PNNs in each channel in isolation (Bankhead et al., 2017). Colabeled PV/PNNs were then quantified by PV/PNN quantification overlap. Cell number was normalized to the area measured in QuPath.

### 2.9 Microglia reconstruction imaging and 3-D analysis

Images of microglia were acquired on a Zeiss 880 Inverted Confocal micropscope. Individual cells were imaged using a 63X magnification with 1.3x optical zoom, capturing the full depth of tissue using a 0.33μm step Z-stack. Images were converted to IMARIS files (v9.5), and “Surfaces” tool was used to render the volume of individual cells, and surface masking was used to reconstruct CD68 within cell volume. Phagocytic capacity was calculated as (total volume CD68 within microglia volume / total volume of that microglia)*100. Individual cell reconstructions were then analyzed using the Sholl statistic setting in Imaris. All reconstructions were performed blinded to experimental condition. At least 4 images were captured per animal, and at least 4 cells/animal were included in all analyses. All statistics were run on biological replicates, which were comprised of the average of all cells for that animal.

### 2.10 P4 endotoxin challenge

To administer LPS or saline at postnatal day 4, pups were injected one at a time to minimize time away from the dam. Pups were briefly removed from the nest and weighed, then placed on a clean towel, where they were injected subcutaneously with 330μg/kg LPS dissolved in 0.9% sterile saline, or 0.9% sterile saline, with a 30-gauge needle to minimize leaking. Tails were then marked with a sharpie before returning to the nest to ensure the same pup was not injected more than once. The total duration of maternal separation for each pup was between 30 seconds to 1 minute. All mice within one litter received the same treatment to prevent indirect exposure to the alternate drug treatment and to avoid differential maternal care. The timing and dose of injection was selected based on previous work from the lab (Bilbo et al., 2005b, 2005a; Bland et al., 2010; Schwarz and Bilbo, 2011; Smith et al., 2020; Williamson et al., 2011).

Following P4 challenge, mice were weaned with littermates into at P27 and transferred to the DID light cycle room at least 2 weeks prior to experiment start. Mice were single housed 1 week prior to DID and DID was performed as described above. This experiment was performed across two separate cohorts which were combined for data analysis. Values between the two cohorts were equally dispersed, indicating low cohort effects on drinking.

### 2.11 2-D Sholl Analysis

2-D Sholl analysis was performed as previously described (Devlin et al., 2025). In brief, immunohistochemistry was performed as above using Iba1 and Tmem119. Images of microglia in the prefrontal cortex were acquired using an Olympus FV3000 inverted confocal at 20X magnification using 5 x 1μM step Z-stacks, then maximum intensity projected. Iba1 and Tmem119 channels were merged, then individual microglia were selected for analysis by a blinded investigator. These cells were pixel segmented using Ilastik software to binarize them (Berg et al., 2019), then run through a custom python script to skeletonize then measure intersections across concentric rings from the soma (Devlin et al., 2025). Images from 4 separate hemispheres per animal were captured, and at least 5 microglia per 20x image were analyzed. Data is presented as all cells analyzed averaged per animal.

### 2.12 Adolescent Drinking in the Dark

The same procedures were followed as in adult DID cohorts, with a few modifications. Due to early age of DID start, mice were not single housed for 1 week prior to drinking. Instead, all mice were weaned into single-housing conditions at P26, and Day 1 of the first week of DID procedures occurred between P28-P31 for all mice. Mice were weighed on Day 1 and Day 3 of drinking each week, and Days 1-2 of drinking were normalized to Day 1 weight, and Days 3-4 of drinking were normalized to Day 3 drinking, in order to account for more rapid changes in weight as mice continued to grow in adolescence. Mice underwent only 2 weeks of DID procedure in order to restrict drinking to the adolescent period of development.

## 3. Results

### 3.1 Constitutive MyD88 removal from microglia does not impact adult voluntary drinking in the DID binge-drinking paradigm

Our first experimental goal was to test the hypothesis that MyD88 loss from microglia would increase voluntary ethanol consumption. To quantify drinking behavior, we used a binge-drinking paradigm called “Drinking in the Dark” (DID). This well-established murine drinking model allows mice to freely consume 20% ethanol four times a week, three days for 2 hours, then for 4 hours on the fourth day, considered the “binge” day (Thiele et al., 2014; Thiele and Navarro, 2014) (Figure 1A). The other three days of the week, the mice are abstinent from alcohol, with free access to water. Alcohol exposure is provided 3 hours into the lights-out part of the mouse 12-12 light cycle, as this period has been identified as the time when mice consume the maximum amount of alcohol (Rhodes et al., 2005). In this model, it has been previously demonstrated that C57BL/6J mice reliably consume 20% alcohol to achieve blood alcohol concentrations (BACs) of at least 1.0 mg/mL without interventions such as food/water restriction or sucrose tapering (Rhodes et al., 2005). The highest average BACs (∼1.6 mg/mL) are observed on the “binge” day, when there are 4 hours of alcohol access. Notably, this paradigm differs from two-bottle choice drinking in the dark (2BC-DID), which allows access to both a 20% alcohol bottle and a water bottle simultaneously for 3 hours. Changes in voluntary consumption were previously observed in whole-body MyD88 knockout male mice in the 2BC-DID method, but the classic one-bottle DID paradigm was not tested (Blednov et al., 2017).

**Figure 1.**
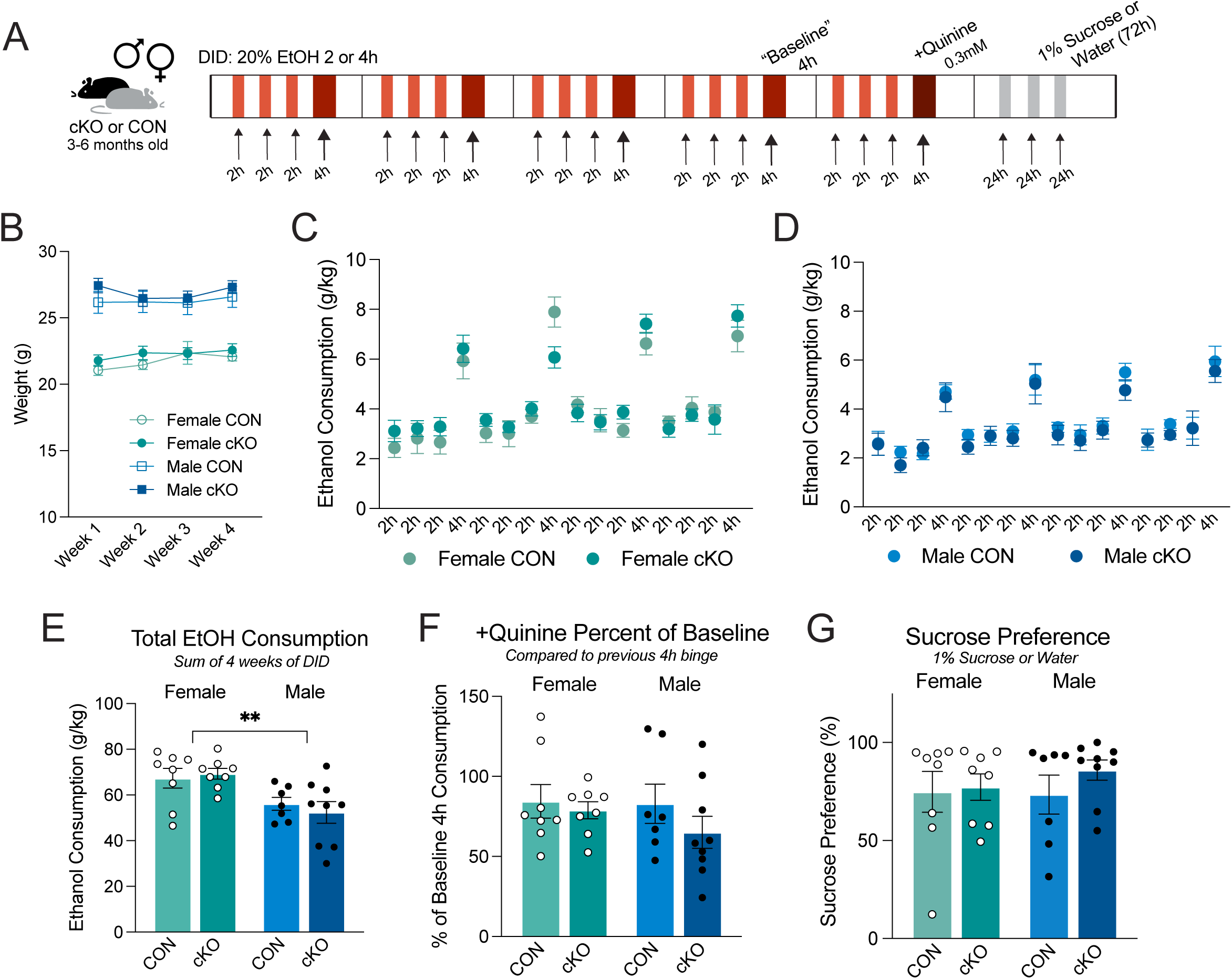
No impact of constitutive microglial-MyD88 removal on voluntary drinking in a one-bottle, binge-like paradigm. (A) Experimental timeline for adult drinking in the dark (DID) experiment. Male and female MyD88-intact (CON) or knockout (cKO) mice were provided with 2 or 4 hours of access to 20% ethanol for 4 weeks, then underwent a quinine challenge in week 5, and a sucrose preference test in week 6. (B) CON and cKO mice weights across drinking exposure. (C) Daily ethanol consumption, normalized to individual weekly body weights, in female CON and cKO mice. (D) Daily ethanol consumption, normalized to individual weekly body weights, in male CON and cKO mice. (E) Total ethanol consumption over 4 weeks of drinking. (F) Impact of quinine bitterant on a 4-hour ethanol binge drinking period, normalized to previous 4-hour consumption. (G) Preference for sucrose water over normal drinking water over a 72-hour period of exposure. All data shown as mean ± SEM. In B-D, 1 point = average of all animals for that experimental group. In E-G, 1 point = average of 1 individual animal.

To quantify consumption on an individual animal level in the one-bottle DID paradigm, male and female adult mice lacking MyD88 in microglia (cKO) and control littermates with intact MyD88 (CON) were single housed for approximately 7 days prior to the first alcohol exposure, as well as for the duration of the experiment. While single housing mice can be considered a stressor, quantification of mouse weight across the 4 weeks of drinking remained consistent (Figure 1B), suggesting that no individual group was significantly more impacted by single housing than others.

Both male and female mice gradually increased their drinking between the first and second week of DID, regardless of genotype (Figure 1C-D). However, there were no significant differences in consumption between genotypes across the 4 weeks of DID (Figure 1C-D), though 2-way mixed effects ANOVA matched by day revealed a trending interaction between genotype x drinking day in females [F_int(15,209)_=1.603, p=0.0749]. When the total consumption across 16 drinking sessions was summed for each individual animal, we observed a significant main effect of sex by 2-way ANOVA, where females drink significantly more than males, consistent with the literature [F_SEX(1,28)_=13.31, p=0.0011] (Figure 1E). However, again, there was no impact of microglial MyD88 genotype on consumption. Together, these first results reveal that in naïve mice, loss of MyD88 from microglia does not affect adult voluntary drinking behaviors in a DID paradigm in either sex.

It has been shown that C57BL/6 mice can exhibit aversion-resistant drinking following alcohol exposure (Lei et al., 2016; Sneddon et al., 2019). This is thought to model a key component of alcohol use disorder: continuing to consume alcohol in the face of negative consequences. In a mouse, this can be modeled through the addition of bitterants to alcohol to determine if mice will continue to drink despite the bad taste. We next sought to determine if MyD88-cKO mice, which do not drink more than CONs at baseline, would exhibit more aversion-resistant drinking than CONs. Drinking more ethanol with quinine would indicate dependence on or desire for alcohol to a greater degree. To test this, we used a 4h binge following initial alcohol exposure (Figure 1A) as their “baseline” consumption. One week later, instead of the normal 20% ethanol in water, the mice received 20% ethanol with 0.3mM quinine. When we normalized the ethanol + quinine consumption to baseline ethanol consumption over a normal 4h binge period, there were no significant differences between groups by 2-way ANOVA, though the majority of mice drank less ethanol + quinine, as expected (Figure 1F). The lack of significant impacts of genotype on drinking when quinine bitterant is present suggests that adults with and without microglial MyD88 exhibit similar levels of aversion-resistant voluntary drinking. Finally, in order to confirm that the loss of MyD88 in microglia does not generally impact reward-seeking behaviors or taste, we administered a sucrose preference test following DID (Figure 1A). Mice were allowed access to both water and 1% sucrose in water for 72 hours. Consumption from both bottles was measured every 24 hours and then averaged across the 3 days of drinking. We found no significant differences in the preference for sucrose across genotype and sex by 2-way ANOVA, with the majority of mice exhibiting a very strong (>80%) preference for sucrose over unsweetened water (Figure 1G).

### 3.2 Adult MyD88-cKO mice do not drink more alcohol across 10 weeks of voluntary alcohol consumption

We next ran a longer experiment to determine if more extended access to voluntary alcohol would reveal differences between CON and cKO mice in drinking behavior. In this next experiment, mice were exposed to 6 weeks of DID, were abstinent for 1 week, then allowed 1 additional 4h binge session. Following these initial 7 weeks, we added a different voluntary consumption paradigm, intermittent 2-bottle choice (2BC). For the following 3 weeks, the same mice underwent 2BC, where for 24 hours, mice had unlimited access to both water in one bottle, and 20% ethanol in the other (Figure 2A). At the end of each 24-hour period, both bottles were measured to determine the preference for alcohol vs. water. Critically, in this paradigm alcohol bottles were removed for 24-48h at a time, to avoid a reduction in consumption that has been observed when mice are given unlimited access.

**Figure 2.**
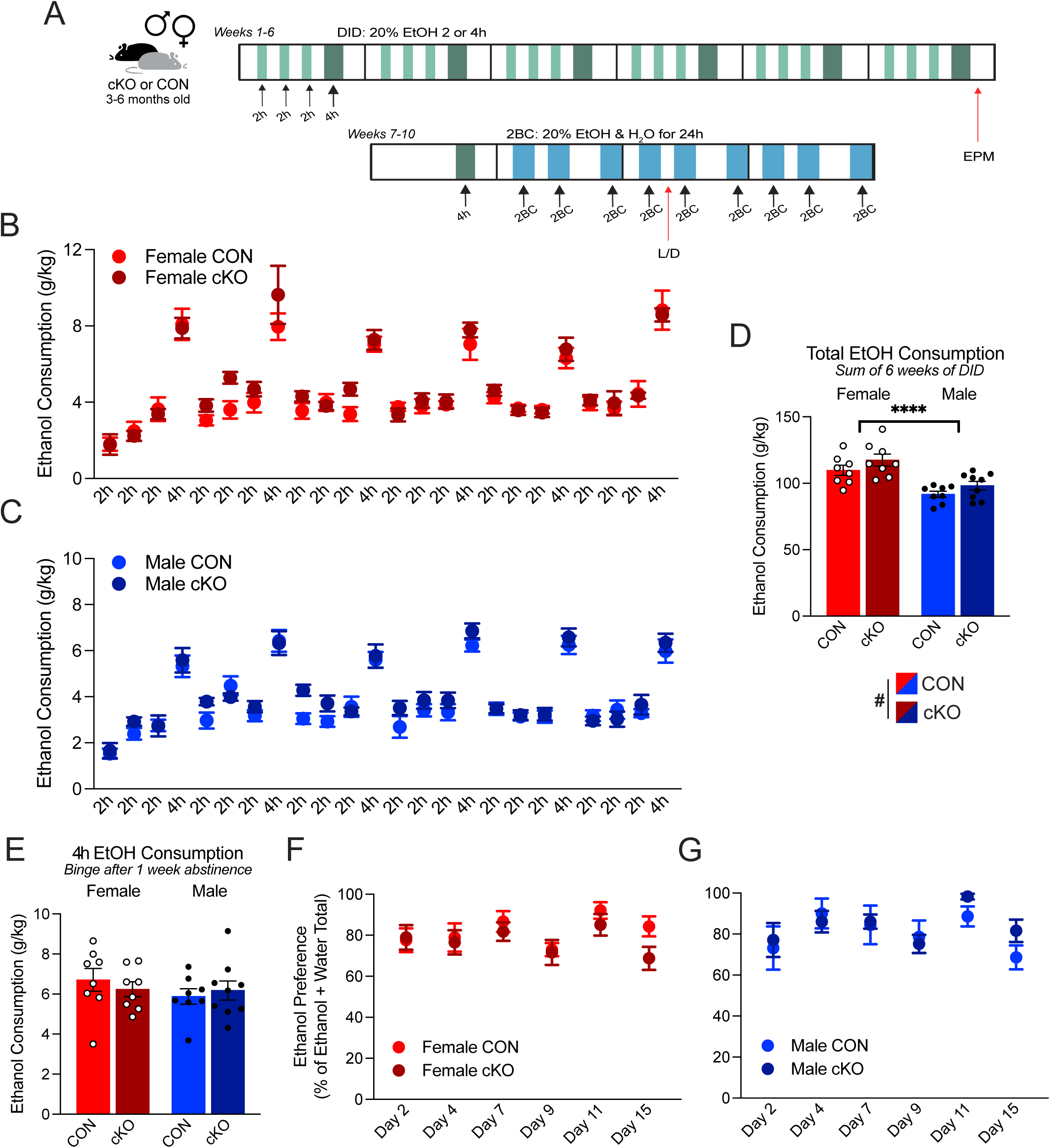
Voluntary drinking in either a binge-like or intermittent access drinking paradigm is not impacted by microglial-MyD88. (A) Experimental paradigm for 10-week DID and 2-bottle choice (2BC) experiment. (B-C) Ethanol consumption normalized to individual weight across 6 weeks of DID in females (B) and males (C). (D) Sum of consumption across 6 weeks of drinking. (E) Total consumption during 4-hour binge period following 7 days of abstinence. (F-G) Ethanol preference over normal drinking water across 3 weeks of 2 bottle choice (2BC) intermittent access paradigm in females (F) and males (G). All data shown as mean ± SEM. In B-C, F-G, 1 point = average of all animals for that experimental group. In D-E, 1 point = average of 1 individual animal.

Neither male nor female CONs or CKOs showed dramatic escalation of consumption across the 6 weeks of DID by 2-way RM ANOVA (Figure 2B-C). However, when the sum of total alcohol consumed over 6 weeks was quantified, we observed a trending (p = 0.05) main effect of genotype by 2-way ANOVA [F_GENO(1,29)_=4.00, p=0.0549], where cKO mice drank more alcohol than CONs (Figure 2D), in support of our original hypothesis. Again, female mice drank more than males across DID, as observed by a significant main effect of sex by 2-way ANOVA [F_SEX(1,28)_=27.46, p<0.0001] (Figure 2D). While not statistically significant, the genotype findings hint that microglial MyD88-dependent signaling may have a minor impact on adult baseline drinking over a six-week period. Thus, they may require either longer access or a challenge to push the immune system to elicit significant changes in drinking.

It has been shown that brief periods of abstinence in mice that are dependent on alcohol lead to greater consumption values upon re-introduction (Becker and Lopez, 2004). If cKO mice drink more than CONs, it could be the case that their dependence grew more rapidly. Therefore, following 6 weeks of DID, we did not provide alcohol for 7 days, and then provided 4h of alcohol access at their normal “binge” time. We did not observe any changes in drinking across any of our groups during this drinking bout by 2-way ANOVA (Figure 2E). At the beginning of the following week, we transitioned to 2BC alcohol administration, allowing mice access for longer than DID (24h at a time). We therefore hypothesized that this extended access would result in greater changes in consumption between our genotypes. However, again we did not see significant differences either between CON and cKO total preference for ethanol across this drinking period, or any drastic changes in preference over time by 2-way RM ANOVA (Figure 2F-G). The preference for ethanol consumption over water did remain high across all groups (between ∼70-90%), confirmed to us that when given a choice, our mice do prefer drinking alcohol, and their consumption during the short bouts of access during DID are not simply due to novelty.

### 3.3 Chronic voluntary alcohol consumption does not change anxiety-like behaviors in MyD88-cKO or CON mice

In the same cohort, we included a water control group, where mice were single housed and provided the same experimental protocol, but with bottles replaced with water instead of 20% ethanol during the binge sessions. This control group allowed for us to determine any impacts of alcohol alone on behavior and brain outcomes in CON mice. We ran two behavioral assays, elevated plus maze (EPM) and light-dark box (LD). One was performed at the conclusion of 6 weeks of DID, and the other during 2BC (Figure 2A). Both EPM and LD are well-established behavioral tests to quantify anxiety-like behaviors in rodents. Previous reports have indicated that chronic alcohol consumption, or withdrawal, can lead to anxiety-like deficits, though there are conflicting reports on whether or not voluntary consumption paradigms allow for enough consumption to induce negative behavioral effects (Salazar and Centanni, 2024). We next wondered if, even though MyD88-cKO and CON mice drink similar volumes of alcohol, one genotype would be more behaviorally impacted by alcohol consumption. For elevated plus maze, mice are placed on a X-shaped apparatus, where one pair of arms are open, without walls, and the other two arms are “closed,” with high walls providing shelter (Figure 3A). Rodents naturally prefer the closed arms but will venture out into the open arms of the maze. Therefore, we can compare the duration of time spent out on the open arms during a 5-minute trial as a proxy for anxiety, with a low duration indicating more “anxious” behavior. We found that neither alcohol nor genotype had any significant impact on EPM behaviors, such as total seconds spent on the open arms of the maze, in males or females by 2-way ANOVA (Figure 3B-C). When we tested anxiety-like behavior with a similar paradigm, light-dark box (Figure 3D), we again observed no changes in any of our groups in either sex by 2-way ANOVA (Figure 3E-F). These experiments suggest that the quantity of alcohol being voluntarily consumed is likely not enough to elicit dramatic changes in behavior, and the loss of microglial-MyD88 in this drinking context does not change these behaviors.

**Figure 3.**
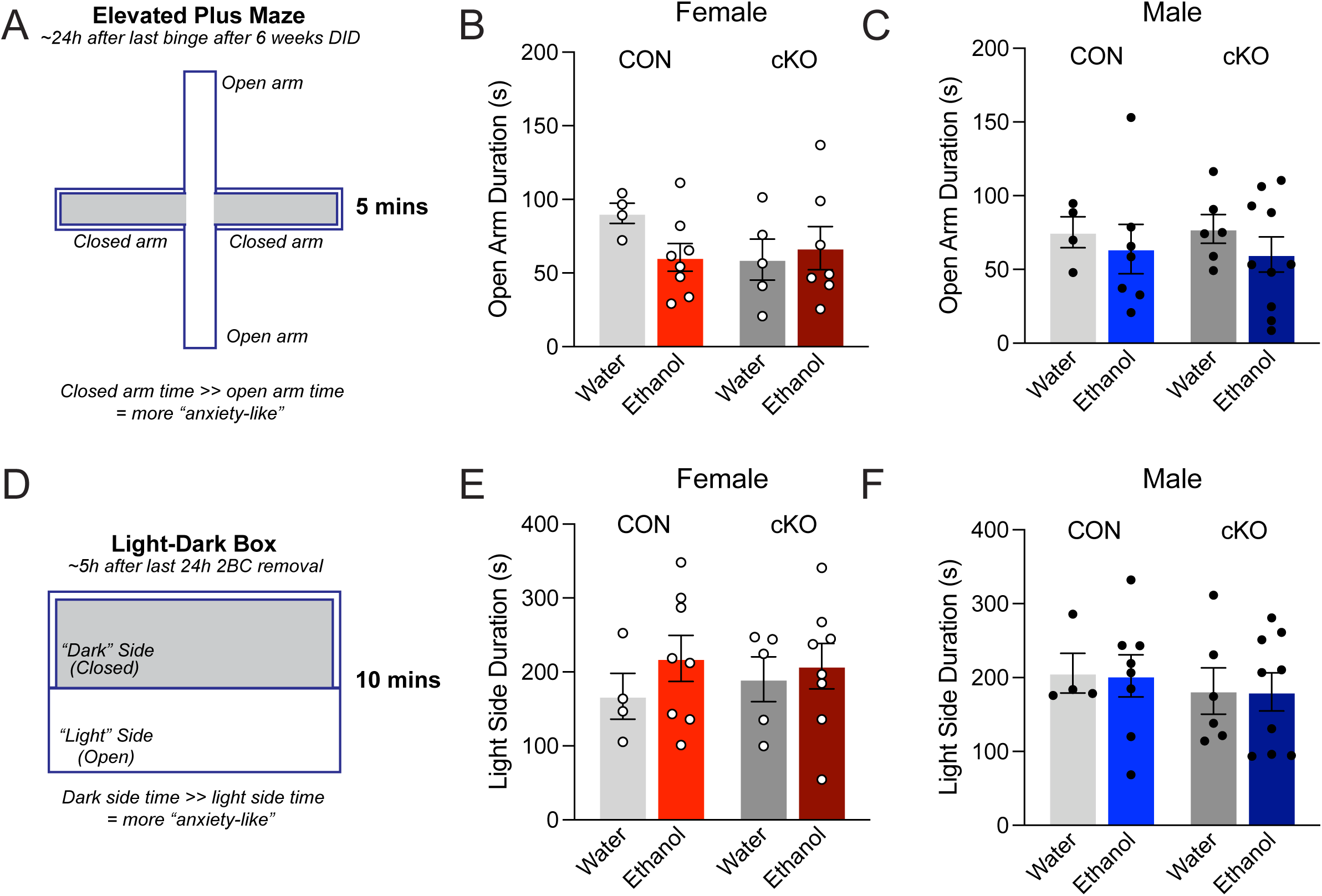
No impacts of MyD88 or alcohol exposure on anxiety-like behaviors in males or females. (A) Diagram of elevated plus maze (EPM) anxiety-like behavior assay. (B-C) Female (B) and male (C) average time in seconds spent in the open arms of the EPM across a 5-minute trial. (D) Diagram of the light-dark box anxiety-like behavior assay. (E-F) Female (E) and male (F) average duration spent on the light side of the apparatus in seconds across a 10-minute trial. All data shown as mean ± SEM, and 1 point = average of 1 individual animal.

### 3.4 Sex and region differences in chronic alcohol consumption and genotype induced impacts on PV cells with PNNs

While we did not observe differences in behavior, we posited that an interaction between alcohol consumption and microglial-MyD88 signaling may still be identifiable within the brain. Therefore, we collected brain tissue 16 hours after the last 24-hour access during 2BC, after 10 weeks of total ethanol (or water) exposure. While each individual mouse drank a different amount of alcohol than any other mouse, the non-significant differences in average consumption between genotypes led us to believe that the brains of these mice can still be grouped and compared.

Previous work in the microglial-MyD88 deficient mice identified that loss of this signaling pathway impacts hippocampal interneuron development (Dziabis et al., 2025). Interestingly, parvalbumin-expressing interneurons (PVIs) are a potential cell target of interest in AUD. PVIs are fast-spiking, GABAergic cells that tightly coordinate the firing of large populations of excitatory neurons to regulate complex, higher-order functions, such as cognition and emotion, which are known to be disrupted both following early life stress and in AUD (Goodwill et al., 2018; Grassi-Oliveira et al., 2016; Ohta et al., 2020). Some of the most extensive, well supported literature on environmental risk factors for AUD demonstrates the strong influence of early life stress on future alcohol and drug abuse in both preclinical and human populations <u>(Enoch, 2011; Loudermilk et al., 2018)</u>. PVIs are highly impacted by inflammation and have a high stress susceptibility (Ruden et al., 2021). Thus, investigating changes in PVIs and their functions from alcohol consumption and the roles MyD88 genotype may play on these cells may provide insight into what neurobiological differences contribute to the development of excessive drinking.

PVI function is particularly critical in the frontal cortex, a late-maturing brain region that is implicated in AUD and addiction broadly due to its importance in decision-making, judgment, and other executive functions. Recently, an interneuron dense ensemble was identified as a high blood ethanol content-responsive in the orbito-prefrontal cortex, and modulation of their activity can change binge-drinking behavior in the DID paradigm (Gimenez-Gomez et al., 2025). Further, other recent work showed that specific ablation of PV interneurons in the dorsal striatum of mice was sufficient to prevent compulsive alcohol drinking, again in a DID paradigm (Patton et al., 2021). These studies provide support for the idea that GABAergic interneurons play a critical role in drinking behaviors that can be quantified by DID. There has also been increasing evidence supporting the hypothesis that perineuronal nets (PNNs), specialized extracellular matrix structures that preferentially surround PVIs to support their function, may perform critical functions in substance use disorders such as AUD. PNNs are hypothesized to play a key role in the stabilization and storage of long-term memories (Lev-Ram et al., 2024; Tsien, 2013). Alcohol use has been shown to increase PNN deposition in brain regions such as the insular cortex and orbitofrontal cortex (Chen et al., 2015; Coleman et al., 2014), but in other brain regions, such as the striatum, chronic drinking leads to decreases in PNNs as well as reduced GABAergic synapses and signaling (Patton et al., 2025). In the insular cortex, the increased PNN presence is critical for aversion-resistant drinking behavior, a hallmark of rodent dependence, as well as a human characteristic of AUD (drinking despite negative consequences) (Chen and Lasek, 2020).

Given the relevance of the interactions between microglia, interneurons, and the extracellular matrix (ECM) in the context of alcohol consumption, we first opted to quantify the deposition of perineuronal nets (PNNs) on parvalbumin+ (PV) interneurons. Changes in PV/PNN colocalization have been previously reported in different brain regions following alcohol drinking (Aguilar and Lasek, 2024). It is thought that increased PNN deposition contributes to the cycle of addiction by stabilizing and protecting addiction-related synaptic connections (Lasek et al., 2018). Based on the existing alcohol literature, as well as what we know about the role of MyD88 signaling in ECM remodeling in the hippocampus (Dziabis et al., 2025), we chose to look at the proportion of PV cells with PNNs in the prefrontal cortex (PFC), the dorsal striatum, and the hippocampus (Figure 4A).

**Figure 4.**
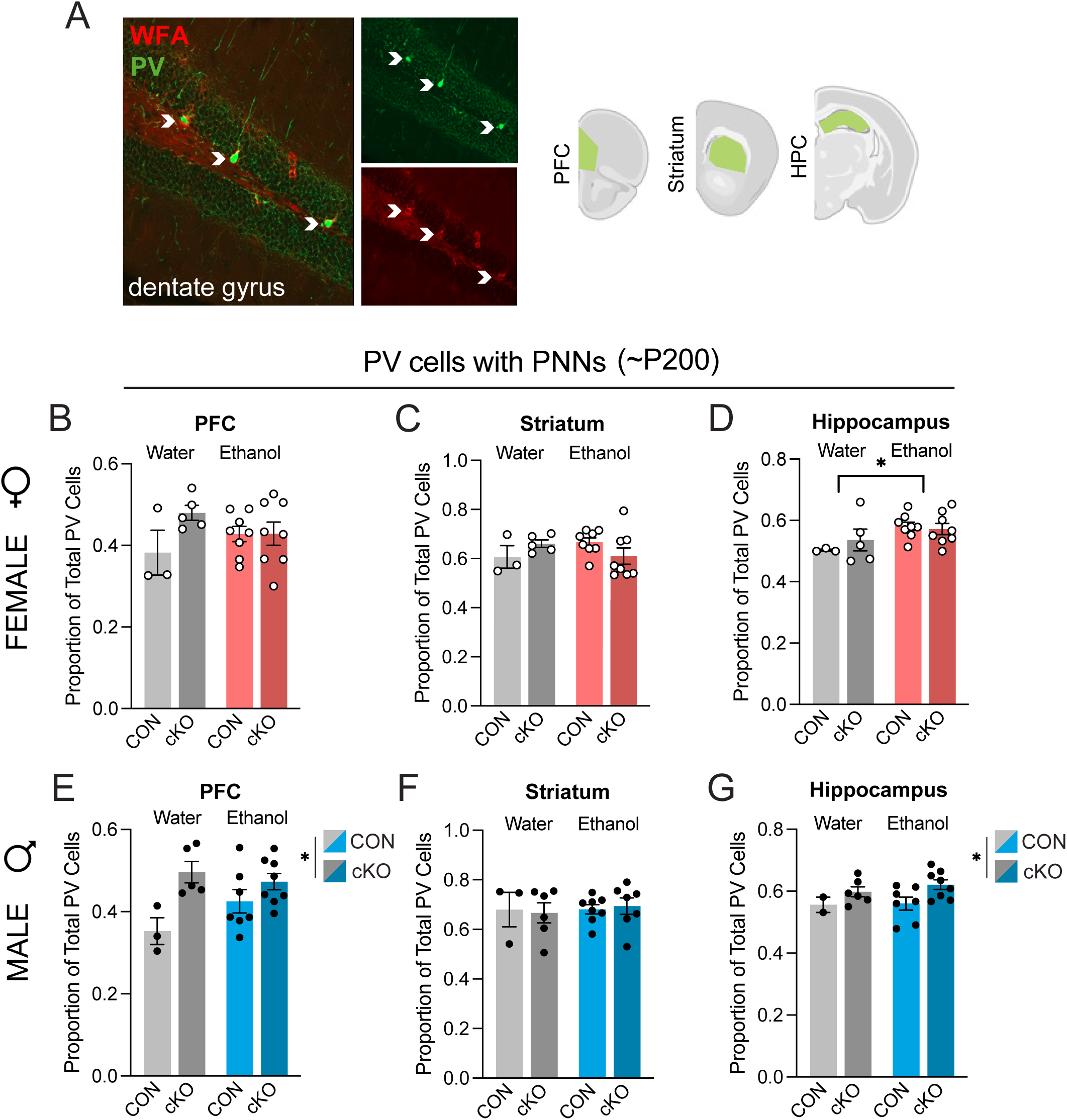
Quantification of perineuronal net deposition following 10 weeks of voluntary alcohol exposure. (A) Representative images of hippocampal parvalbumin interneurons (PV, green) and perineuronal nets (WFA, red), with colocalization indicated by white arrows, and representative areas where colocalization was quantified. (B-D) Quantification of proportion of PV interneurons with PNNs surrounding them in the prefrontal cortex (PFC, B), striatum (C) and dorsal hippocampus (D) in females. (E-G). Quantification of proportion of PV interneurons with PNNs surrounding them in the prefrontal cortex (PFC, E), striatum (F) and dorsal hippocampus (G) in males. All data shown as mean ± SEM, and 1 point = average of at least 2 images for 1 individual animal.

Across our three brain regions of interest, in females, we did not observe any significant genotype-dependent changes by 2-way ANOVA (Figure 4B-D). However, there was a main effect of alcohol consumption on the proportion of PV cells with PNNs in the hippocampus, where there was a greater amount of PNN+ PV cells [F_ALC(1,20)_=6.028, p=0.0234] (Figure 4D). In males, we found no changes in the striatum (Figure 4F), but in the PFC and the hippocampus, there were significant main effects of genotype on PNN+ PV cells by 2-way ANOVA. In both the PFC [F_GENO(1,19)_=11.39, p=0.0032] (Figure 4E) and the hippocampus [F_GENO(1,19)_=5.164, p=0.0349] (Figure 4G), cKO mice had a higher proportion of PV cells with PNNs around them. While the functional consequence of this finding is not known, based on previous work in the field, one might expect that more ECM deposited about PV cells is likely working to stabilize synapses, and perhaps to reinforce behaviors related to drinking. Previous work from our lab and others have identified the importance of microglial signaling and phagocytic activity in the regulation and remodeling of ECM in brain regions such as the hippocampus (Dziabis et al., 2025; Nguyen et al., 2020). Therefore, we next wondered if alcohol or MyD88-cKO impacted microglia in these same brain regions.

### 3.5 Minimal impacts on male and female microglia morphology following 10 weeks of voluntary drinking

Microglia have been previously shown to be extensively impacted by alcohol, even at the level at voluntary consumption (McCarthy et al., 2018). Alcohol is an inflammatory agent and can cause extensive damage in the brain and body. Alcohol itself can cross the blood-brain barrier and may act directly on microglial receptors. Alcohol also leads to the production of reactive oxygen species (ROS) and immune molecules like cytokines, resulting in microglial activation. This has been shown both in preclinical work, as well as in postmortem studies of individuals who suffered from AUD (He and Crews, 2008; Qin and Crews, 2012b). We hypothesized that MyD88-deficient microglia would be less responsive to the inflammatory effects of alcohol consumption, and that this lack of “inflammatory” response in the brain would lead to less aversive experiences, and therefore more drinking.

One hallmark of microglial “activation” is changes in morphology. The shape of microglia does not necessarily correspond to a specific function (Paolicelli et al., 2022), so if changes in morphology are observed, they often require further investigation in order to draw conclusions about microglial response. Microglia that are actively responding to an acute inflammatory event often pull in their processes, become bushy, and upregulated CD68+ lysosomes, which can be further investigated to confirm if this is indicative of increased phagocytic activity. We sought to determine if alcohol-induced changes in microglial volume or lysosomal content were modulated by loss of microglial MyD88 in males or females. We volumetrically reconstructed microglia and their lysosomes in the same three brain regions (PFC, striatum, and hippocampus). Overall, there were no changes in microglia volume in any group, male or female – 10 weeks of alcohol exposure did not change the average microglia volume (data not shown). In females, we observed no changes in lysosomal volume per cell in any of the three regions by 2-way ANOVA (Figure 5A-C). In males, there was a trending main effect by 2-way ANOVA of alcohol treatment on lysosomal volume in the striatum, where ethanol microglia contained slightly more CD68% [F_ALC(1,23)_=3.757, p=0.065] (Figure 5E). Interestingly, there was also a nearly significant interaction effect on PFC lysosomal content [F_int(1,22)_=3.949, p=0.0595] (Figure 5D). This seems to be mostly driven by the cKO-alcohol group, where the animals are stratified into a high-content and low-content grouping. This heterogeneity is reminiscent of previous work in our lab that showed a highly heterogeneous response of male offspring frontal cortex microglia to prenatal challenge (Block et al., 2022). When we correlated average lysosomal content in PFC microglia from individual animals to their average alcohol consumption, we found no significant correlation, suggesting this stratification in the cKO alcohol male PFC microglia is likely not driven by differences in drinking.

**Figure 5.**
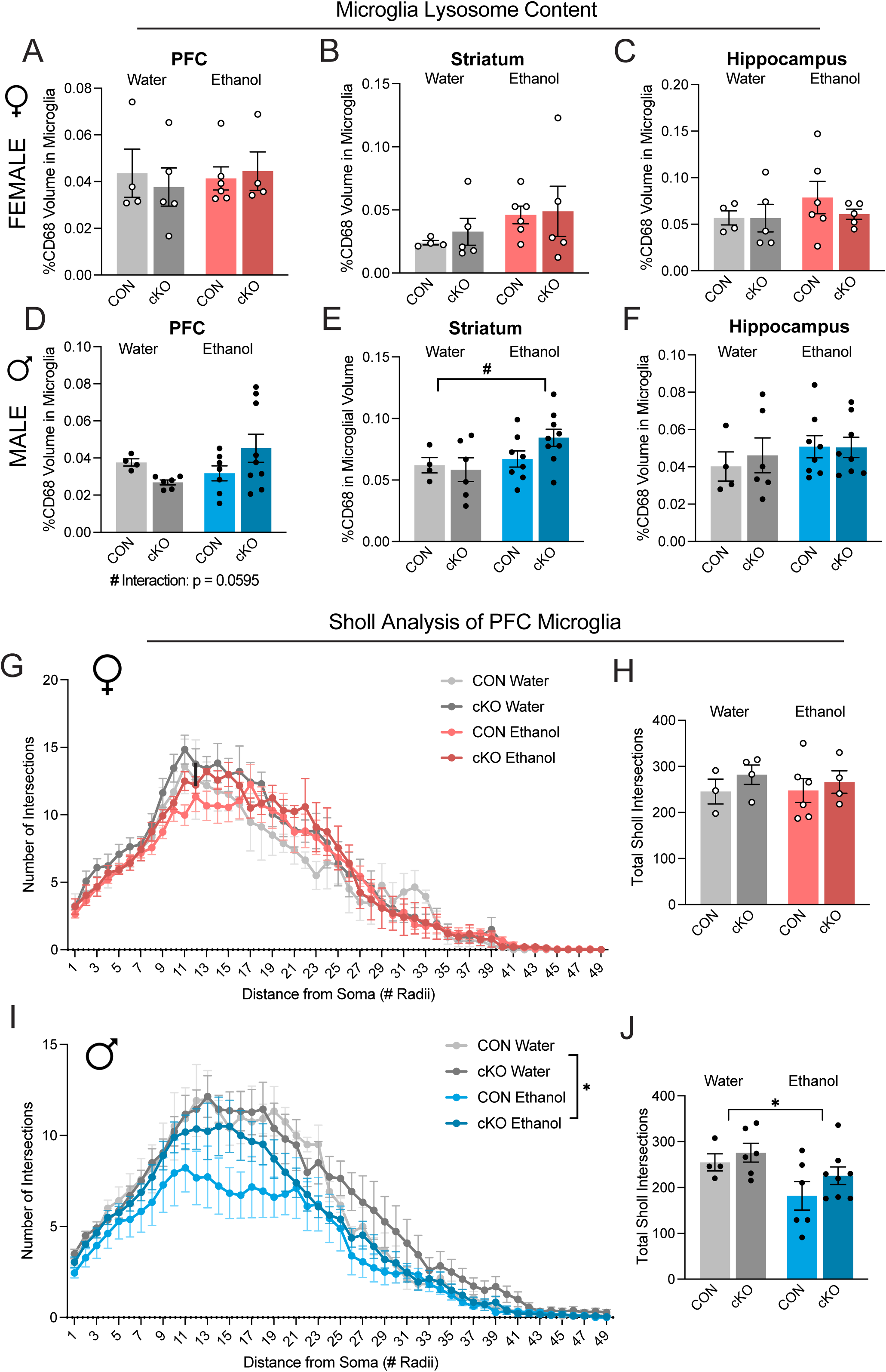
Broad morphological characterization of microglia following 10 weeks of voluntary alcohol exposure. (A-C) Quantification of lysosome (CD68+) volume normalized to individual microglia volume across the PFC (A), striatum (B), and hippocampus (C) in females. (D-F) Quantification of lysosome (CD68+) volume normalized to individual microglia volume across the PFC (D), striatum (E), and hippocampus (F) in males. (G-J) Sholl intersections quantification of microglial morphology in the PFC of females (G-H) and males (I-J). All data shown as mean ± SEM. For A-F, H, and J, 1 point = average of at least 4 microglia for 1 individual animal.

Next, we wanted to determine how chronic drinking impacts microglia through Sholl analysis, to measure the impact of alcohol on ramification. Here, we focused on the microglia in the PFC, where we observed both the most genotype-induced changes in PNN deposition and interesting lysosomal changes in males. We found that the morphology of female PFC microglia was not impacted by alcohol or genotype by 3-way RM ANOVA [F_ALC(1,13)_=0.0653, p=0.8022; F_GENO(1,13)_=1.079,p=0.3178] (Figure 5G), and the quantification of total intersections was unchanged by 2-way ANOVA (Figure 5H). Male microglia, on the other hand, saw ramification changes as a result of alcohol exposure by both 3-way RM ANOVA (Figure 5I) and 2-way ANOVA of total interactions [F_ALC(1,20)_=6.499, p=0.0191] (Figure 5J). While this impact of ethanol was not significantly rescued by MyD88 removal, the average of the alcohol-cKO animals falls between water and alcohol-CON values, supporting our hypothesis that MyD88-cKO microglia are semi-protected from alcohol-induced morphology changes, though only in males.

### 3.6 Early life experience impacts adult voluntary alcohol consumption in males only

As previously shown, loss of MyD88 from microglia alone does not impact the brain enough to elicit large changes in voluntary alcohol consumption in a one-bottle DID paradigm. We therefore sought to “push” our model through an additional inflammatory event prior to drinking exposure. A major risk factor for later-life alcohol abuse is early exposure to drinking alcohol (Dawson et al., 2008), and early life inflammation through events like early life stress have demonstrated impacts on risk for developing AUD <u>(Enoch, 2011; Loudermilk et al., 2018)</u>. It is well established that perturbations to developing neurons and their circuits impact adult behavior. Thus, investigating changes to normal developmental mechanisms may provide insight into what neurobiological differences contribute to the development of excessive drinking.

We wondered if early life exposure to an inflammatory agent that activates the MyD88 signaling pathway would elicit changes in voluntary drinking between CON and cKO mice. To run this experiment, a moderate dose of lipopolysaccharide (LPS) or saline control was administered subcutaneously at postnatal day 4 to CON or cKO mice. In young adulthood (2-3 months), these mice were exposed to 3 weeks of DID (Figure 6A). When we looked at female drinking, we found little impact of P4 LPS treatment or genotype on daily consumption values by mixed-effects ANOVA, matched by day of consumption [F_TX(1,17)_=1.162, p=0.2961; F_GENO(1,17)_=1.415, p=0.02505] (Figure 6B) or the sum of total consumption across 3 weeks of DID by 2-way ANOVA [F_TX(1,13)_=0.1929, p=0.6677; F_GENO(1,13)_=0.2676, p=0.1676] (Figure 6C). When we ran a mixed-effects ANOVA on male daily drinking (matched by day), we observed significant main effects of treatment [F_TX(1,35)_=17.68, p=0.0002], genotype [F_GENO(1,35)_=6.008, p=0.0194], and a genotype x treatment significant interaction effect [F_int(1,35)_=8.168, p=0.0071] (Figure 6D). When we quantified male total consumption, we observed a main effect of treatment [F_TX(1,33)_=16.18, p=0.0003] and a genotype x treatment interaction effect [F_int(1,33)_=8.49, p=0.0063] by 2-way ANOVA (Figure 6E). Bonferroni post-hoc found that all groups drank significantly more than saline-CONs (Figure 6E), in almost a stepwise manner, where saline-cKO drank second most. (p=0.0169), then LPS treated CON (p=0.0003) and cKO mice (p=0.0009) both drank even more, all compared to saline-CONs. However, when we ran 2-D Sholl analysis on male PFC microglia from saline- and LPS-injected CON and cKO mice that underwent 3 weeks of DID, we did not observe significant changes in ramification from either treatment or genotype (Supplemental Figure 1A-B). While male microglial ramification was previously differentially impacted by ethanol in MyD88-CONs and cKOs (Figure 5I-J), these changes were observed after ∼10 weeks of drinking, suggesting that morphological changes to microglia may require longer alcohol exposure to occur. So, while LPS treatment in CON males clearly increased total consumption compared to saline-injected CONs, both LPS- and saline-injected male cKO groups exhibited increased drinking later in life. This suggests that the impact of this manipulation cannot be entirely attributed to the specific inflammatory agent being injected. These data reveal a male-specific vulnerability to early life challenge-induced changes in adult voluntary drinking.

**Figure 6.**
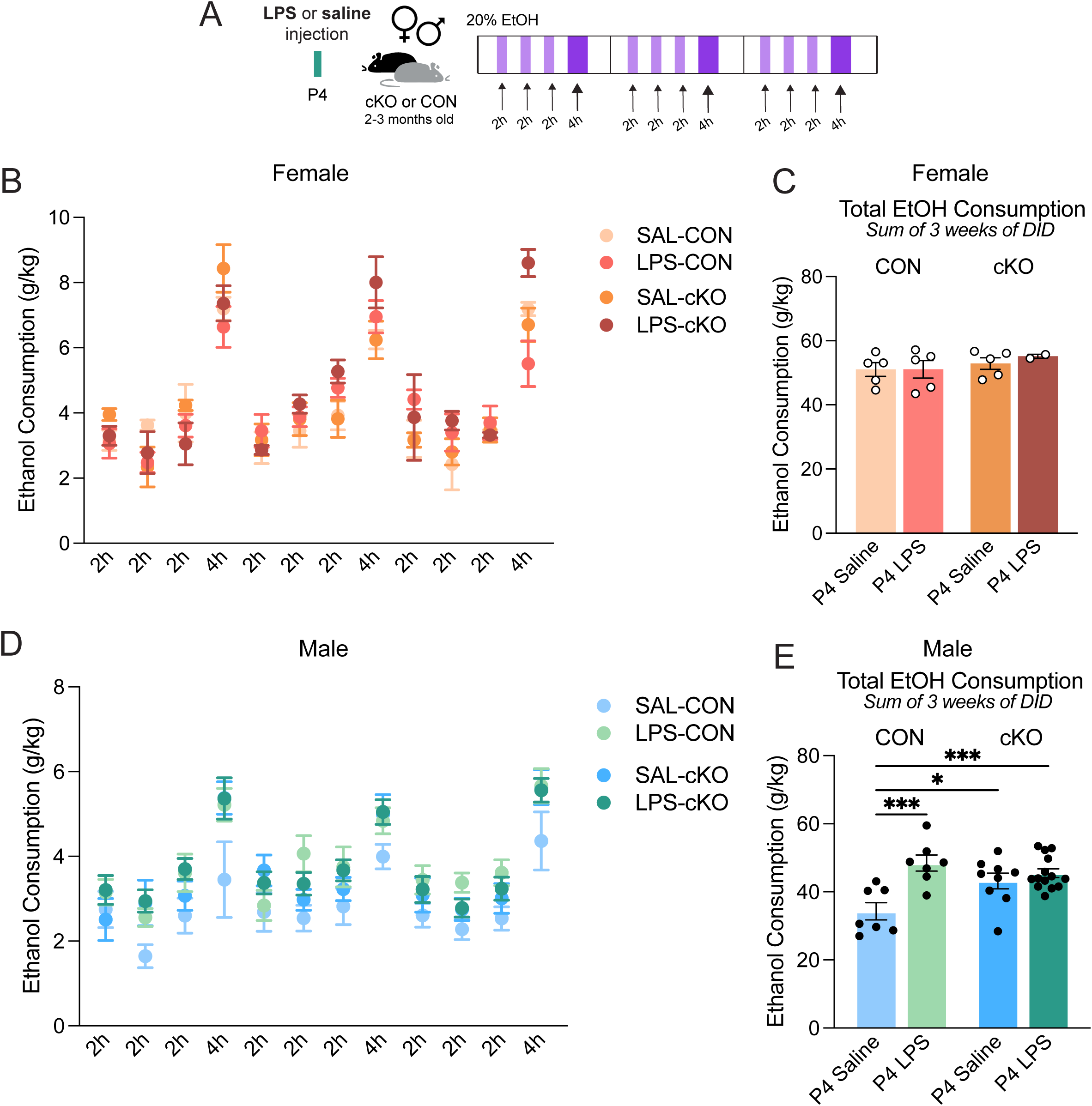
Impact of early life injection on adult drinking behavior in MyD88-CON and cKO mice. (A) Experimental timeline for early life endotoxin experiment. Male and female MyD88 CON and cKO mice were injected at postnatal day (P)4 with either lipopolysaccharide (LPS) endotoxin or saline control. In adulthood, mice were then exposed to the DID binge-like voluntary drinking paradigm for 3 weeks. (B) Daily consumption of ethanol by female CON and cKO mice. (C) Sum of total consumption over 3 weeks of DID in females. (D) Daily consumption of ethanol by male CON and cKO mice. (E) Sum of total consumption over 3 weeks of DID in males. All data shown as mean ± SEM. In B and D, 1 point = average of all animals for that experimental group. In C and E, 1 point = average of 1 individual animal.

## 4. Discussion

Here, we show that the relationship between microglial MyD88 signaling and voluntary alcohol consumption in adulthood is highly context dependent. In our hands, adult microglial MyD88 deletion had minimal effects on male and female voluntary drinking behavior in a one-bottle, binge-drinking paradigm (DID) or in an intermittent access two-bottle paradigm (2BC). However, alcohol intake in this model likely did not reach levels sufficient to induce negative affect anxiety-like behaviors typically observed in dependence models of AUD, such as vapor exposure. Similarly, alcohol exposure did not impact MyD88 loss-induced increases in perineuronal net (PNN) deposition in the male hippocampus and PFC. However, alcohol exposure did lead to changes in male microglia ramification by Sholl analysis, independent of MyD88 genotype. Interestingly, early life injection, whether of bacterial endotoxin (in MyD88-CON mice and MyD88-cKO mice) or of saline control (in MyD88-cKO mice), had the potential to unmask a male-specific increase in adult alcohol drinking, indicating that developmental perturbations can be critical determinants of later-life alcohol consumption behaviors.

Neuroimmune signaling has been implicated in each phase of the cycle of addiction, and microglia are increasingly recognized as central mediators of alcohol-induced brain changes. Clinical studies consistently report significantly elevated peripheral circulating cytokines in individuals with AUD, in particular IL-6, IL-7, IL-8, and TNF, during both active drinking and withdrawal (Adams et al., 2020). This is also true in the brain; human postmortem tissue, mostly from frontal cortex tissue, consistently show that AUD patients exhibit large gene expression changes compared to matched control tissue, many of which are relevant to immune signaling (Flatscher-Bader et al., 2005; Lewohl et al., 2000; Liu et al., 2006; Mayfield et al., 2002). Looking at protein endpoints, microglial morphology changes classically associated with inflammation are repeatedly reported via histology, as well as upregulation of TLRs by western blot (Crews et al., 2013; He and Crews, 2008). However, whether immune changes are primary drivers of brain changes in AUD, or secondary consequences of the inflammatory effects of excessive alcohol consumption, is still unresolved.

TLR4, which signals in part through MyD88, has been proposed to play an outsized role in direct responses to alcohol (<u>Fernandez-Lizarbe et al., 2009</u>; Holloway et al., 2023). Global deletion of TLR4 or MyD88 blunted the brain’s canonical immune response to alcohol, including the upregulation of cytokines and chemokines (Holloway et al., 2023). TLR4 can be activated by damage-associated molecular patterns, such as HMBG1, an endogenous stress molecule that is released following alcohol exposure, as well as by circulating endotoxin from alcohol-induced gut permeability (Bishehsari et al., 2017; Crews et al., 2024). Our findings here build on this foundation, demonstrating that cell-specific disruption of MyD88 in microglia does not greatly alter voluntary drinking behavior in adulthood in a traditional DID paradigm, unless preceded by an early life manipulation, highlighting the importance of developmental context in shaping later alcohol sensitivity.

Innate immune signaling molecules such as cytokines are of critical importance to both brain development and adult brain functioning. Cytokines influence synaptic function, learning, and memory, particularly in the hippocampus, and chronic inflammation can disrupt the balance needed for cognitive flexibility and executive control (Ferro et al., 2021). Chronic inflammation is also a known contributor to PV dysfunction (Ruden et al., 2021), as well as to remodeling of the extracellular matrix (ECM), including PNNs, with consequences for addiction-relevant behaviors (Chaunsali et al., 2021; Fawcett et al., 2019; Lasek et al., 2018). Consistent with the literature, we observed increased PNN deposition in male MyD88-cKO mice, suggesting that microglial innate immune signaling contributes to ECM regulation in regions strongly implicated in alcohol use, such as the PFC.

PNNs are classically thought to restrict plasticity, and their removal/degradation is associated with the reopening of plasticity windows. This experience-dependent remodeling is mediated in part through ECM cleavage proteases, such as matrix metalloproteases and ADAMTS family members, many of which are released by microglia (Lasek et al., 2018). Microglia also remodel ECM through phagocytic mechanisms in the hippocampus, with consequences for learning and memory (Nguyen et al., 2020). These findings position microglia as key regulators of PNN dynamics and suggest a potential mechanism through which early immune perturbation and MyD88 signaling could bias circuits in addiction-relevant brain regions.

Our findings are consistent with recent work using the same microglial MyD88 conditional knockout line in forced alcohol exposure paradigms. In a high-dose injection model, microglia undergo rapid and sustained morphological changes, increased synaptic engulfment behavior, and ECM-related transcriptional signatures following alcohol exposure. Deletion of MyD88 from microglia (in both Cx3cr1-BT-Cre and Tmem119-Cre lines) was sufficient to rescue these changes (Paouri et al., 2025). Notably, these effects were observed under conditions of severe alcohol exposure, which induce sedation and neuronal stress, suggesting that microglial MyD88 signaling may be most relevant in high-consumption or dependence-associated states. This interpretation aligns with studies using microglial depletion tools, such as CSF1R antagonist PLX, which demonstrate that microglia contribute to escalated drinking and anxiety-like behaviors only after dependence induction by vapor exposure, not during non-dependent drinking, such as what we are modeling with DID (Warden et al., 2020b). We did not observe significant escalation in drinking across multiple weeks of DID, and normal laboratory inbred strains of mice have been reported to not voluntarily drink to dependence, or the levels induced in Paouri et al. (Barkley-Levenson and Crabbe, 2014). However, because we did not explicitly measure BACs in these experiments, this interpretation can only be speculative.

There are several important limitations to this specific work. First, despite the lack of significant differences observed in adult drinking between CON and cKO mice in the 1 bottle DID paradigm, the role of microglial MyD88 in drinking behaviors cannot be conclusively determined here. Voluntary drinking is highly context-dependent, and how much mice drink will vary significantly based on the access paradigm and environmental setting. The existing literature that supports a role for MyD88 signaling on voluntary drinking utilized a number of different drinking paradigms, including varied alcohol concentrations, where the authors consistently observed increased drinking in male whole-body MyD88 knockout mice, and no significant changes in drinking in female whole-body knockouts (Blednov et al., 2017). These paradigms include a variation on traditional one-bottle DID, where there are 2 bottles (one with 20% ethanol, the other with water) during the drinking periods, a 2-bottle choice, every-other-day intermittent access paradigm (with 15% then 20% ethanol), and a 2-bottle choice continuous access paradigm. There were no changes in either sex with continuous access reported in this study, which is consistent with the literature on the importance of intermittent access for alcohol consumption (Blednov et al., 2017). Critically, while the primary paradigm we used in this study, one-bottle DID, was measured in other transgenic lines in Blednov et al., the MyD88 KO line was not tested using this paradigm. Without testing drinking in our microglial MyD88-cKO mouse using two-bottle DID, our ability to compare between these experiments is limited. Due to the context sensitivity of drinking behaviors and the confounds of differences across institutional environments, interpretation of this work would benefit from both a reproduction of the full-body MyD88 KO effect in our facility prior to more definitively determining the role of the MyD88-cKO on drinking behaviors. Further, the specific ages of the mice used for experiments reported in Blednov et al. is somewhat limited (> 2 months of age) (Blednov et al., 2017). Our early-life challenge cohort was assessed for alcohol consumption behavior changes at a younger age to reduce the interval of time between neonatal manipulations and drinking (2-3 months old), while the other adult cohorts were assessed at a slightly older age (3-6 months old). We ran additional experiments where the one-bottle DID paradigm began immediately after weaning (between P28-P31), during adolescence (Supplemental Figure 2A). We again observed no changes in voluntary drinking behavior between genotypes in males or females (Supplemental Figure 2B-D), suggesting a young age of drinking onset does not influence this behavior acutely. While both the 2-3- and 3-6-month cohorts are considered adult ages, there is a lack of studies directly comparing voluntary drinking changes across this age gap, and therefore age-dependent effects on drinking cannot be entirely excluded. Future studies examining the persistence of this neonatal injection phenotype into later adult age ranges or repeating the one-bottle DID experiments closer to 2 months of age will be necessary to know if the effects observed are stable over a range of adult ages.

One additional caveat of these findings is the transgenic mouse line selected to test the hypothesis regarding the role of microglial MyD88 in alcohol-related behaviors. While our lab has extensively validated the removal of MyD88 from microglia in the brain over a variety of methods (Dziabis et al., 2025; Rawls et al., 2025; Rivera et al., 2019; Smith et al., 2020), this removal was not directly validated again in these experiments. Perhaps of more importance, however, is that microglia are not the only cells in the body that express Cx3cr1, and there are likely peripheral macrophages that are impacted by MyD88 loss. Because alcohol is orally consumed, other Cx3cr1-expressing cells, such as tissue-resident macrophages, like those in the gut, may be contributing to our findings. Therefore, this work may be better interpreted as the loss of MyD88 across the broader Cx3cr1-expressing lineage of cells was not sufficient to induce changes in drinking in the DID paradigm. This is still important, as other critical cell types implicated in alcohol behaviors, such as neurons and astrocytes, also express MyD88, and should not be impacted in this transgenic line. Additionally, recent work from another research group using the same Cx3cr1-BT-Cre crossed with MyD88-f/f from our colony in a different alcohol paradigm successfully replicated their findings using a new, more microglia-specific Cre driver, Tmem119-Cre, crossed with MyD88-f/f animals (Paouri et al., 2025). Validation of our findings in this mouse line would help elucidate the roles of microglia vs. peripheral monocyte MyD88 signaling in drinking behavior. Additionally, future studies incorporating histological and molecular analyses of microglia in early life challenge cohorts will be necessary to determine whether persistent alterations in microglial state may be contributing to the later life behavioral phenotypes observed.

Together, these data suggest that microglial MyD88 signaling may play a limited role in the initiation of naïve, adult alcohol consumption, but this role may be better detected depending on early life experience. Our initial hypothesis was that early-life inflammation through an endotoxin exposure may lower the threshold for engaging these microglial-dependent mechanisms in a sex-specific manner, therefore increasing vulnerability to excessive drinking later in life. We were surprised to find that a moderate dose of LPS at postnatal day 4 was sufficient to change young adult drinking behavior in CON males, which may be related to “immune priming” of neonatal microglia to later life challenges, a phenomena that has been repeatedly reported in the context of exposure to immune stimuli (Bilbo and Schwarz, 2009; Carloni et al., 2021). However, the observation that even saline-injected MyD88-cKO males exhibited elevated drinking relative to controls highlights the potential sensitivity of developmental windows to very subtle perturbations, including the mild stress of neonatal handling, which has been previously shown to produce lasting effects on stress circuitry and behavior (Liu et al., 1997), including through microglia themselves (Carloni et al., 2021; Melbourne et al., 2021). The surprising finding that the injection alone, in absence of an inflammatory stimulus, was sufficient to “unmask” genotypic changes at a level comparable to an LPS challenge in a control animal highlights the critical importance of subtle changes in early life experience or context on later-life behaviors, particularly in genetically predisposed organisms.

Finally, the clinical relevance of targeting neuroimmune pathways in AUD is underscored by the emerging efficacy of apremilast, a phosphodiesterase type 4 (PDE4) inhibitor, initially approved by the FDA for use in humans to treat psoriasis. Apremilast has been shown to reduce both alcohol consumption behaviors across multiple strains of mice with high excessive drinking risk and also to decrease drinking in a clinical heavy-drinking population (Grigsby et al., 2023). While the precise neural mechanisms underlying these effects are currently still under investigation, emerging work has pointed to an interaction between apremilast and GABAergic transmission in the context of drinking (Blednov et al., 2024; Vozella et al., 2025). Our findings support the idea that modulation of microglial immune signaling, particularly in those with heightened stress or inflammatory histories or genetic predispositions, may represent a promising therapeutic strategy.

## Credit authorship contribution statement

**Julia E. Dziabis**: Conceptualization, Methodology, Validation, Formal analysis, Investigation, Data curation, Project administration, Visualization, Writing – original draft, Writing – review & editing. **Neil Rogers**: Methodology, Software, Validation, Formal analysis, Investigation, Data curation, Visualization. **Benjamin L. Horvath**: Methodology, Validation, Formal analysis, Investigation, Data curation, Visualization. **Michael S. Patton**: Methodology, Validation, Investigation. **Irene O. Jonathan**: Investigation, Data curation, Visualization. **Erica J. Freeman**: Investigation. **Will Sun**: Investigation. **Jerome Moulden II**: Investigation. **Grace Zhang**: Investigation. **Staci D. Bilbo**: Conceptualization, Methodology, Recourses, Project administration, Funding acquisition, Writing – review & editing, Supervision.

## Acknowledgements

This work was supported by NIH grants U01-AA029969 (to SDB) and F31-AA030712 (to JED). The authors thank the Duke Light Microscopy Core Facility for microscopy and analysis support, and to the Integrative Neuroscience Initiative on Alcoholism – Neuroimmune for their many valuable insights on this project. The authors have no conflicting financial interests.

---

**Supplemental Figure 1. Male microglial ramification is not changed after 3 weeks of adult drinking following early life injection**

(A-B) 2-D Sholl intersections quantification of microglial morphology in the PFC of males. All data shown as mean ± SEM. For B, 1 point = average of at least 20 microglia for 1 individual animal.

**Supplemental Figure 2. Loss of microglial-MyD88 does not impact adolescent voluntary alcohol consumption in a one-bottle drinking in the dark paradigm**

(A) Experimental timeline for adolescent drinking experiment. Male and female MyD88 CON and cKO mice were weaned into single-housing conditions at postnatal day (P)26, then the first day of DID binge-link voluntary drinking paradigm began on P28-P31. Mice underwent 2 weeks of DID. (B) Daily consumption of ethanol by female CON and cKO adolescent mice. (C) Daily consumption of ethanol by male CON and cKO adolescent mice. (D) Sum of total consumption over 2 weeks of DID in females and males. All data shown as mean ± SEM. In B and C, 1 point = average of all animals for that experimental group for that day of drinking. In D, 1 point = sum of 1 individual animal.

