## Supplementary figures and images for "Context-dependent effects of microglial MyD88 removal on voluntary ethanol consumption in mice"

### Supplemental Figures 1-2

Sholl Analysis of Male P4 Saline or LPS PFC Microglia

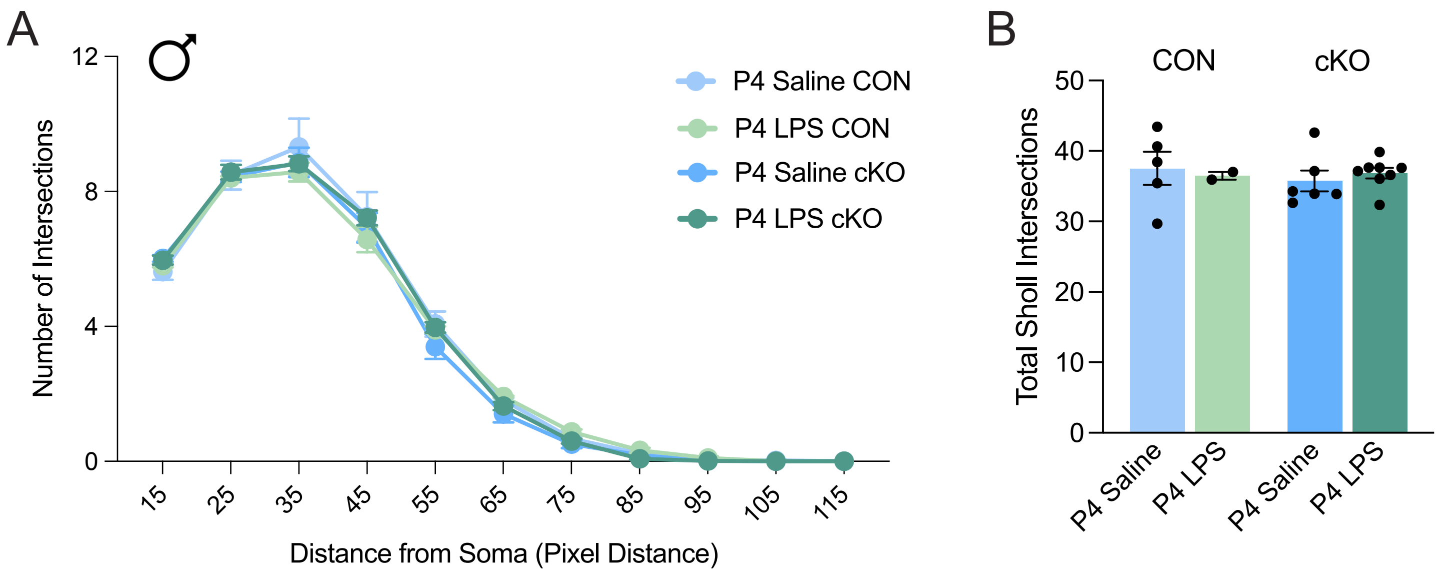

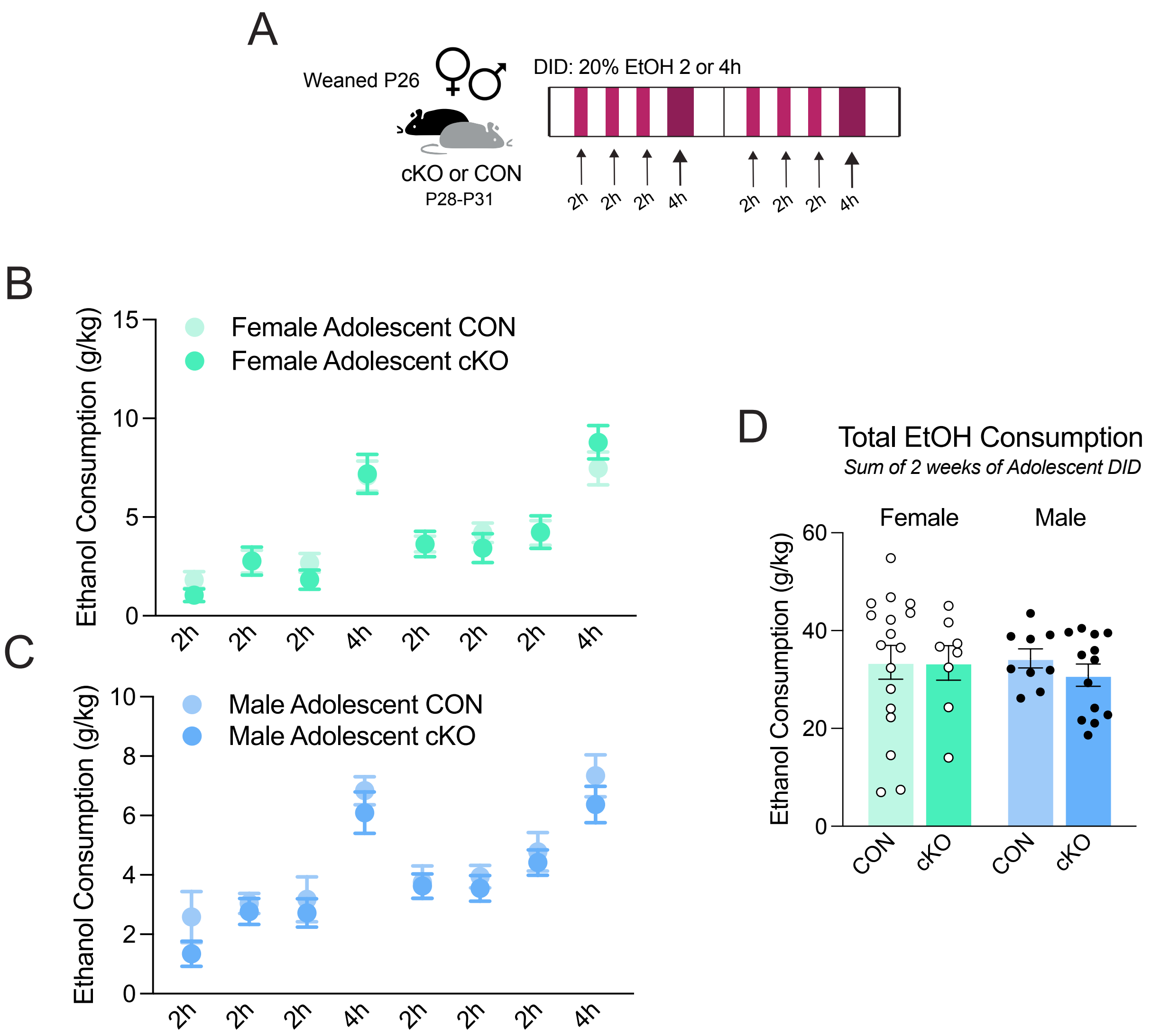
